# An antisense RNA capable of modulating the expression of the tumor suppressor microRNA-34a

**DOI:** 10.1101/234310

**Authors:** Jason T. Serviss, Felix Clemens Richter, Jimmy Van den Eynden, Nathanael Andrews, Miranda Houtman, Mattias Vesterlund, Laura Schwarzmueller, Per Johnsson, Erik Larsson, Dan Grandér, Katja Pokrovskaja Tamm

## Abstract

The microRNA-34a is a well-studied tumor suppressor microRNA (miRNA) and a direct downstream target of TP53 with roles in several pathways associated with oncogenesis, such as proliferation, cellular growth, and differentiation. Due to its broad tumor suppressive activity, it is not surprising that *miR34a* expression is altered in a wide variety of solid tumors and hematological malignancies. However, the mechanisms by which *miR34a* is regulated in these cancers is largely unknown. In this study, we find that a long non-coding RNA transcribed antisense to the *miR34a* host gene, is critical for *miR34a* expression and mediation of its cellular functions in multiple types of human cancer. We name this long non-coding RNA *lncTAM34a,* and characterize its ability to facilitate *miR34a* expression under different types of cellular stress in both *TP53* deficient and wild type settings.

## Introduction

In recent years advances in functional genomics have revolutionized our understanding of the human genome. Evidence now points to the fact that approximately 75% of the genome is transcribed but only ~1.2% of this is responsible for encoding proteins (International Human Genome Sequencing Consortium 2004, Djebali et al. 2012). Of these recently identified elements, long non-coding (lnc) RNAs are defined as transcripts exceeding 200 base pairs (bp) in length with a lack of a functional open reading frame. Some lncRNAs are dually classified as antisense (as) RNAs that are expressed from the same locus as a sense transcript in the opposite orientation. Current estimates using high-throughput transcriptome sequencing, indicate that up to 20-40% of the approximately 20,000 protein-coding genes exhibit antisense transcription (Chen et al. 2004, Katayama et al. 2005, Ozsolak et al. 2010). Systematic large-scale studies have shown aberrant expression of asRNAs to be associated with tumorigenesis (Balbin et al. 2015) and, although characterization of several of these has identified asRNA-mediated regulation of multiple well known tumorigenic factors (Yap et al. 2010, Johnsson et al. 2013), the vast majority of potential tumor-associated lncRNAs have not yet been characterized. The known mechanisms by which asRNAs accomplish their regulatory functions are diverse, and include recruitment of chromatin modifying factors (Rinn et al. 2007, Johnsson et al. 2013), acting as microRNA (miRNA) sponges (Memczak et al. 2013), and causing transcriptional interference (Conley et al. 2012).

Responses to cellular stress, e.g. DNA damage, sustained oncogene expression, and nutrient deprivation, are all tightly controlled cellular pathways that are almost universally dysregulated in cancer. Cellular signaling, in response to these types of stresses, often converges on the transcription factor TP53 that regulates transcription of coding and non-coding downstream targets. One important non-coding target of TP53 is the tumor suppressor miRNA known as *miR34a* (Raver-Shapira et al. 2007). Upon TP53 activation *miR34a* expression is increased allowing it to down-regulate target genes involved in cellular pathways such as growth factor signaling, apoptosis, differentiation, and cellular senescence (Lal et al. 2011, Slabakova et al. 2017). Thus, *miR34a* is a crucial factor in mediating activated TP53 response and, the fact that it is often deleted or down-regulated in human cancers indicates, its tumor suppressive effect and makes it a valuable prognostic marker (Cole et al. 2008, Gallardo et al. 2009, Zenz et al. 2009, Cheng et al. 2010, Liu et al. 2011). Reduced *miR34a* transcription is mediated via epigenetic regulation in many solid tumors, including colorectal-,pancreatic-, and ovarian cancer (Vogt et al. 2011), as well as numerous types of hematological malignancies (Chim et al. 2010). In addition, *miR34a* has been shown to be transcriptionally regulated via TP53 homologs, TP63 and TP73, other transcription factors, e.g. STAT3 and MYC, and, in addition, post-transcriptionally through miRNA sponging by the NEAT1 lncRNA (Chang et al. 2008, Su et al. 2010, Agostini et al. 2011, Rokavec et al. 2015, Ding et al. 2017). Despite these findings, the mechanisms underlying *miR34a* regulation in the context of oncogenesis have not yet been fully elucidated.

Studies across multiple cancer types have reported a decrease in oncogenic phenotypes when *miR34a* expression is induced in a TP53-null background, although endogenous mechanisms for achieving this have not yet been discovered (Liu et al. 2011, Ahn et al. 2012, Yang et al. 2012, Stahlhut et al. 2015, Wang et al. 2015). In addition, previous reports from large-scale studies interrogating global TP53-mediated regulation of lncRNAs have identified a lncRNA (known as RP3-510D11.2 and LINC01759) originating in the antisense orientation from the *miR34a* locus that is induced upon numerous forms of cellular stress (Rashi-Elkeles et al. 2014, Hunten et al. 2015, Leveille et al. 2015, Ashouri et al. 2016, Kim et al. 2017). Despite this, none of these studies have functionally characterized this transcript, which we name *L*ong-*N*on-*C*oding *T*ranscriptional *A*ctivator of *MiR34a* (lncTAM34a). In this study we functionally characterize the *lncTAM34a* transcript, and find that it positively regulates *miR34a* expression resulting in a decrease of several tumorigenic phenotypes. Furthermore, we find that *lncTAM34a-mediated* up-regulation of *miR34a* is sufficient to induce endogenous cellular mechanisms counteracting several types of stress stimuli in a TP53-deficient background. Finally, similar to the functional roles of antisense transcription at protein-coding genes, we identify a rare example of an antisense RNA capable of regulating a cancer-associated miRNA.

## Results

### *lncTAM34a* is a broadly expressed non-coding transcript whose levels correlate with *miR34a* expression

*lncTAM34a* is transcribed in a “head-to-head” orientation with approximately 100 base pair overlap with the *miR34a* host gene (HG) (**Fig. 1a**). Due to the fact that sense/antisense pairs can be both concordantly and discordantly expressed, we sought to evaluate this relationship in the case of *miR34a* HG and its asRNA. Using a diverse panel of cancer cell lines, we detected coexpression of both the *miR34a* HG and *lncTAM34a* (**Fig. 1b**). We used cell lines with a known *TP53* status in the panel due to previous reports that *miR34a* and *lncTAM34a* are known downstream targets of TP53. These results indicate that *miR34a* HG and *lncTAM34a* are co-expressed and that their expression levels correlate with *TP53* status, with *TP53^-/-^*cells tending to have decreased or undetectable expression of both transcripts.

**Figure 1:**
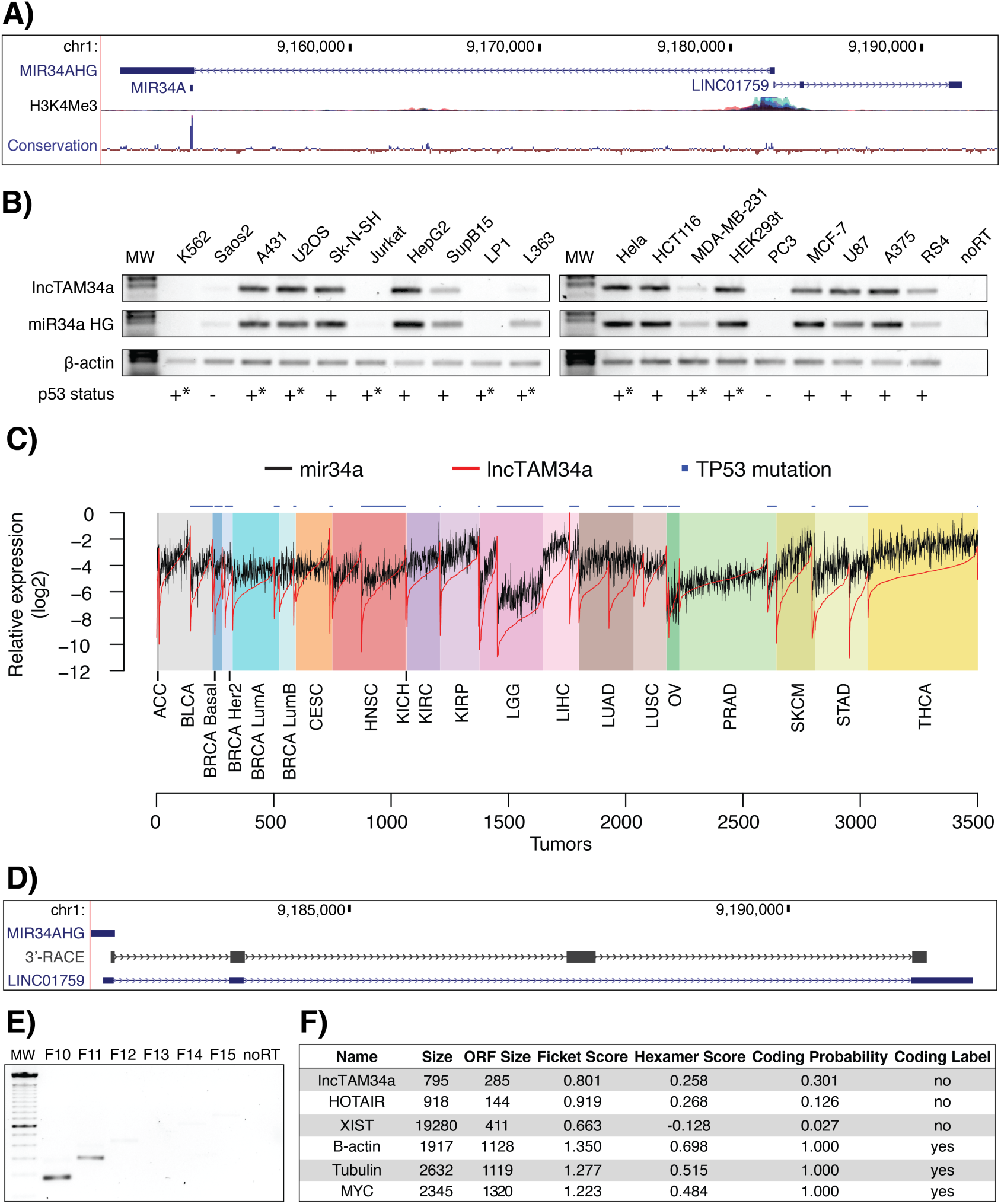
Characterization of the *lncTAM34a* transcript. **A)** Architecture of the *miR34a* locus (hg38, RefSeq) including *miR34a* HG, mature *miR34a*, and *lncTAM34a (LINC01759)*. H3K4me3 ChIP-seq data, indicating the active promoter region, and conservation are also shown. **B)** Semi-quantitative PCR data from the screening of a panel of cancer cell lines. Wild type *TP53* is indicated with +, - indicates null, and +* represents either a non-null *TP53* mutation or wild-type *TP53* with mechanisms present that inhibit its function (e.g. SV40 large T antigen in HEK293T cells). **C)** TCGA correlation analysis. Expression was log2 normalized to the maximum expression value. Nonsynonymous *TP53* mutations are indicated on the top of the plot (cancer type abbreviation definitions and corresponding statistics are in Figure 1-Supplement 1). **D)** 3’-RACE sequencing results and the annotated *lncTAM34a (LINC01759)* are shown. **E)** Semi-quantitative PCR results from the primer walk assay (i.e. common reverse primer (exon 2) and forward primers (F10-F15) staggered upstream of *lncTAM34a’s* annotated start site) performed using HEK293T cells (Figure 1-Supplement 2a details primer placement) **F)** Coding potential analysis assessed via the Coding-potential Assessment Tool including *lncTAM34a*, two known non-coding RNAs (*HOTAIR* and *XIST*), and three protein-coding RNAs (β-actin, Tubulin, and *MYC*).

We next sought to analyze primary cancer samples to examine whether a correlation between *lncTAM34a* and *miR34a* expression levels could be identified. We utilized RNA sequencing data from The Cancer Genome Atlas (TCGA) after stratifying patients by cancer type, *TP53* status, and, in the case of breast cancer, cancer subtypes. The results indicate that *lncTAM34a* and *miR34a* expression are strongly correlated in the vast majority of cancer types examined, both in the presence and absence of wild-type *TP53* (**Fig. 1c, Figure 1-Figure Supplement 1a**). The results also further confirm that the expression levels of both *miR34a* and *lncTAM34a* are significantly reduced in patients with nonsynonymous *TP53* mutations (**Figure 1-Figure Supplement 1b**).

Next, we aimed to gain a thorough understanding of *lncTAM34a*’s molecular characteristics and cellular localization. To experimentally determine the 3’ termination site for the *lncTAM34a* transcript we performed 3’ rapid amplification of cDNA ends (RACE) using the U2OS osteosarcoma cell line that exhibited high endogenous levels of *lncTAM34a* in the cell panel screening. Sequencing the cloned cDNA indicated that the transcripts 3’ transcription termination site is 525 bp upstream of the *lncTAM34a* transcript’s annotated termination site (**Fig. 1d**). Next, we characterized the *lncTAM34a* 5’ transcription start site by carrying out a primer walk assay, i.e. a common reverse primer was placed in exon 2 and forward primers were gradually staggered upstream of *lncTAM34a*’s annotated start site (**Figure 1-Figure Supplement 2a**). Our results indicated that the 5’ start site for *lncTAM34a* is in fact approximately 90 bp (F11 primer) to 220 bp (F12 primer) upstream of the annotated start site (**Fig. 1e**). Polyadenylation status was evaluated via cDNA synthesis with either random nanomers or oligo(DT) primers followed by semi-quantitative PCR which showed that *lncTAM34a* is polyadenylated although the unspliced form seems to only be present in a polyadenylation negative state (**Figure 1-Figure Supplement 2b**). Furthermore, we investigated the propensity of *lncTAM34a* to be alternatively spliced in U2OS cells, using PCR cloning followed by sequencing and found that the transcript is post-transcriptionally spliced to form multiple isoforms (**Figure 1-Figure Supplement 2c**). In order to evaluate the subcellular localization of *lncTAM34a,* we made use of RNA sequencing data from five cancer cell lines included in the ENCODE (Encode Project Consortium 2012) project that had been fractionated into cytosolic and nuclear fractions. The analysis revealed that the *lncTAM34a* transcript primarily localizes to the nucleus with only a minor fraction in the cytosol (**Figure 1-Figure Supplement 2d**).

Lastly, we utilized several approaches to evaluate the coding potential of the *lncTAM34a* transcript. The Coding-Potential Assessment Tool is a bioinformatics-based tool that uses a logistic regression model to evaluate coding-potential by examining open reading frame (ORF) length, ORF coverage, Fickett score, and hexamer score (Wang et al. 2013). Results indicated that *lncTAM34a* has a similar low coding capacity to known non-coding transcripts such as *HOTAIR* and *XIST* (Fig. 1F). We further confirmed these results using the Coding-Potential Calculator that uses a support vector machine-based classifier and accesses an alternate set of discriminatory features (**Figure 1-Figure Supplement 2e**) (Kong et al. 2007). Finally, we downloaded mass spectrometry spectra for 11 cancer cell lines (Geiger et al. 2012), 7 of which were also present in the cell line panel above (**Fig. 1b**), and searched it against a database of human protein sequences which also contained the 6 frame translation of *lncTAM34a*. However, we did not manage to detect any peptides matching the sequence in any of the 11 cell lines. Taken together our results indicate that *lncTAM34a* is not a coding transcript and that it is not translated to any significant degree.

### TP53-mediated regulation of *lncTAM34a* expression

*miR34a* is a known downstream target of TP53 and has been previously shown to exhibit increased expression within multiple contexts of cellular stress. Several global analyses of TP53-regulated lncRNAs have also shown *lncTAM34a* to be induced upon TP53 activation (Rashi-Elkeles et al. 2014, Hunten et al. 2015, Leveille et al. 2015, Ashouri et al. 2016, Kim et al. 2017). To confirm these results in our biological systems, we treated HEK293T, embryonic kidney cells, and HCT116, colorectal cancer cells, with the DNA damaging agent doxorubicin to activate TP53. QPCR-mediated measurements of both *miR34a* HG and *lncTAM34a* indicated that their expression levels were increased in response to doxorubicin treatment in both cell lines (**Fig. 2a**). To assess whether TP53 was responsible for the increase in *lncTAM34a* expression upon DNA damage, we treated *TP53*^+/+^ and *TP53^-/-^* HCT116 cells with increasing concentrations of doxorubicin and monitored the expression of both *miR34a* HG and *lncTAM34a*. We observed a dose-dependent increase in both *miR34a* HG and *lncTAM34a* expression levels with increasing amounts of doxorubicin, revealing that these two transcripts are co-regulated, although, this effect was largely abrogated in *TP53^-/-^*cells (Fig. 2b). These results indicate that TP53 activation increases *lncTAM34a* expression upon DNA damage. Nevertheless, *TP53^-/-^* cells also showed a dose-dependent increase in both *miR34a* HG and *lncTAM34a*, suggesting that additional factors, other than *TP53* are capable of initiating an increase in expression of both of these transcripts upon DNA damage.

**Figure 2:**
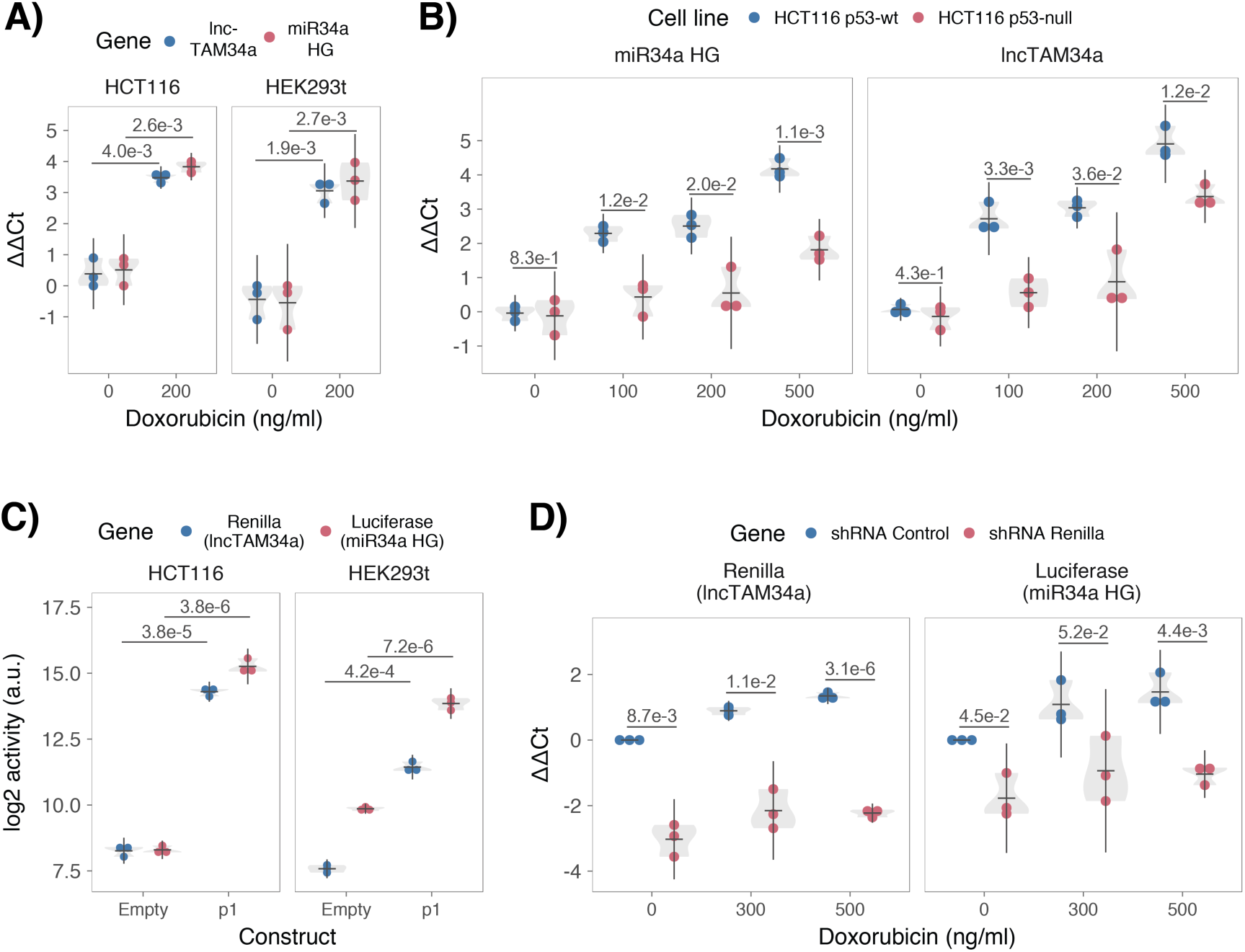
TP53-mediated regulation of the *miR34a* locus. **A)** Evaluating the effects of 24 hours of treatment with 200 ng/ml doxorubicin on *lncTAM34a* and *miR34a* HG in HCT116 and HEK293T cells.* **B)** Monitoring *miR34a* HG and *lncTAM34a* expression levels during 24 hours of doxorubicin treatment in *TP53^+/+^* and *TP53^-/-^* HCT116 cells.* **C)** Quantification of luciferase and renilla levels after transfection of HCT116 and HEK293T cells with the p1 construct (**Figure 2 - Supplement 2** contains a schematic representation of the p1 construct).* **D)** HCT116 cells were co-transfected with the p1 construct and shRNA renilla or shRNA control and subsequently treated with increasing doses of doxorubicin. 24 hours post-treatment, cells were harvested and renilla and luciferase levels were measured using QPCR.* *Individual points represent results from independent experiments and the gray shadow indicates the density of those points. Error bars show the 95% CI, black horizontal lines represent the mean, and *P* values are shown over long horizontal lines indicating the comparison tested. All experiments in Figure 2 were performed in biological triplicate.

The head-to-head orientation of *miR34a* HG and *lncTAM34a*, suggests that transcription is initiated from a single promoter in a bi-directional manner (**Fig 1a**). To investigate whether *miR34a* HG and *lncTAM34a* are transcribed from the same promoter as divergent transcripts, we cloned the previously reported *miR34a* HG promoter, a ~300 bp region including the TP53 binding site and the majority of the first exon of both transcripts, into a luciferase/renilla dual reporter vector (**Figure 2-Figure Supplement 1a-b**) (Raver-Shapira et al. 2007). We hereafter refer to this construct as p1. Upon transfection of p1 into HCT116 and HEK293T cell lines we observed increases in both luciferase and renilla indicating that *miR34a* HG and *lncTAM34a* expression can be regulated by a single promoter contained within the p1 construct (**Fig. 2c**).

### *lncTAM34a* facilitates *miR34a* induction in response to DNA damage

We hypothesized that *lncTAM34a* may regulate *miR34a* HG levels and, in addition, that the overlapping regions of the sense and antisense transcripts may mediate this regulation. Knockdown of endogenous *lncTAM34a* is complicated by its various isoforms (**Figure 1-Figure Supplement 2c**). For this reason, we utilized the p1 construct to evaluate the regulatory role of *lncTAM34a* on *miR34a* HG. Accordingly, we first co-transfected the pi construct, containing the overlapping region of the two transcripts, and two different short hairpin (sh) RNAs targeting renilla into HEK293T cells and subsequently measured luciferase and renilla expression. The results indicated that shRNA-mediated knock-down of the p1-renilla transcript (corresponding to *lncTAM34a*) caused p1-luciferase (corresponding to *miR34a* HG) levels to concomitantly decrease (**Figure 2-Figure Supplement 2**). The results suggest that *lncTAM34a* positively regulates levels of *miR34a* HG and that the transcriptional product of *lncTAM34a* within the pi construct contributes to inducing a *miR34a* response. To further support these conclusions and better understand the role of *lncTAM34a* during TP53 activation, *TP53^+/+^* HCTii6 cells were co-transfected with p1 and shRNA renilla (2.1) and subsequently treated with increasing doses of doxorubicin. Again, the results showed a concomitant reduction in luciferase levels upon knock-down of p1 -renilla i.e. the *lncTAM34a* corresponding segment of the p1 transcript (**Fig. 2d**). Furthermore, the results showed that in the absence of p1-renilla the expected induction of p1-luciferase in response to TP53 activation by DNA damage is abrogated. Collectively these results indicate that *lncTAM34a* positively regulates *miR34a* expression and furthermore, suggests that it is crucial for an appropriate TP53-mediated *miR34a* response to DNA damage.

### *lncTAM34a* can regulate *miR34a* host gene independently of *TP53*

Despite the fact that TP53 regulates *miR34a* HG and *lncTAM34a* expression, our results showed that other factors are also able to regulate this locus (**Fig. 2b**). Utilizing a lentiviral system, we stably over-expressed the *lncTAM34a* transcript in three *TP53*-null cell lines, PC3 (prostate cancer), Saos2 (osteogenic sarcoma), and Skov3 (ovarian adenocarcinoma). We first analyzed the levels of *lncTAM34a* in these stable cell lines, compared to HEK293T cells, which have high endogenous levels of *lncTAM34a*. On average, the over-expression was approximately 30-fold higher in the over-expression cell lines than in HEK293T cells, roughly corresponding to physiologically relevant levels in cells encountering a stress stimulus, such as DNA damage (**Figure 3-Figure Supplement 1**). Analysis of *miR34a* levels in the *lncTAM34a* over-expressing cell lines showed that this over-expression resulted in a concomitant increase in the expression of *miR34a* in all three cell lines (**Fig. 3a**). These results indicate that, in the absence of *TP53, miR34a* expression may be rescued by activating *lncTAM34a* expression.

**Figure 3:**
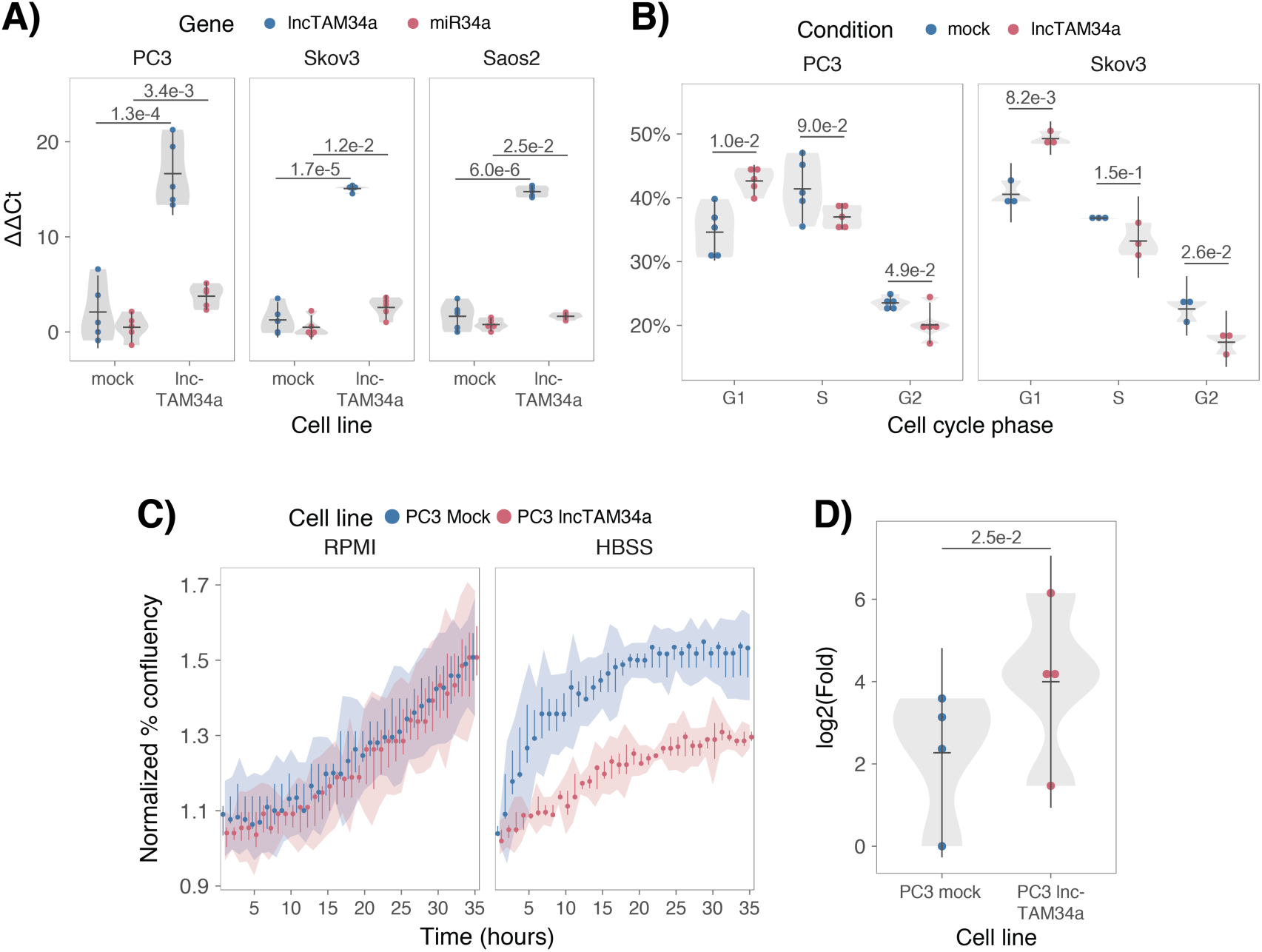
*lncTAM34a* positively regulates *miR34a* and its associated phenotypes. **A)** QPCR-mediated quantification of *miR34a* expression in cell lines stably over-expressing *lncTAM34a*.* **B)** Cell cycle analysis comparing stably over-expressing *lncTAM34a* cell lines to the respective mock control.* **C)** Analysis of cellular growth over time in *lncTAM34a* over-expressing PC3 cells. Points represent the median from 3 independent experiments, the colored shadows indicate the 95% confidence interval, and vertical lines show the minimum and maximum values obtained from the three experiments. **D)** Differential phosphorylated polymerase II binding in *lncTAM34a* over-expressing PC3 cells.* *Individual points represent results from independent experiments and the gray shadow indicates the density of those points. Error bars show the 95% CI, black horizontal lines represent the mean, and *P* values are shown over long horizontal lines indicating the comparison tested.

*miR34a* has been previously shown to regulate cell cycle progression, with *miR34a* induction causing G1 arrest (Raver-Shapira et al. 2007, Tarasov et al. 2007). Cell cycle analysis via determination of DNA content showed a significant increase in G1 phase cells and a concomitant decrease in G2 phase cells in the PC3 and Skov3 *lncTAM34a* over-expressing cell lines, indicating G1 arrest (**Fig. 3b**). The effects of *miR34a* on the cell cycle are mediated by its ability to target cell cycle regulators such as cyclin D1 *(CCND1) (Sun et al. 2008)*. Quantification of both *CCND1* RNA expression (**Figure 3-Figure Supplement 2a**) and protein levels (**Figure 3-Figure Supplement 2b**) in the PC3 *lncTAM34a* over-expressing cell line showed a significant decrease of *CCND1* levels compared to the mock control. Collectively, these results indicate that *lncTAM34a*-mediated induction of *miR34a* is sufficient to result in the corresponding miR34a-directed effects on cell cycle.

*miR34a* is also a well-known inhibitor of cellular growth via its ability to negatively regulate growth factor signaling. Furthermore, starvation has been shown to induce *miR34a* expression causing inactivation of numerous prosurvival growth factors (*Lal et al. 2011*). We further interrogated the effects of *lncTAM34a* over-expression by monitoring the growth of the PC3 stable cell lines in both normal and starvation conditions via confluency measurements over a 35-hour period. Under normal growth conditions there is a small but significant reduction (*P* = 3.0e-8; linear regression, **Fig. 3c**) in confluency in the *lncTAM34a* over-expressing cell lines compared to mock control. However, these effects on cell growth are drastically increased in starvation conditions (*P* = 9.5e-67; linear regression; **Fig. 3c**). This is in agreement with our previous results, and suggests that *lncTAM34a*-mediated increases in *miR34a* expression are crucial under conditions of stress and necessary for the initiation of an appropriate cellular response. In summary, we find that over-expression of *lncTAM34a* is sufficient to increase *miR34a* expression and gives rise to known phenotypes observed upon induction of *miR34a*.

### *lncTAM34a* transcriptionally activates *miR34a* host gene

Antisense RNAs have been reported to mediate their effects both via transcriptional and post-transcriptional mechanisms. Due to the fact that *miR34a* expression is undetected in wild type PC3 cells (**Fig. 1b**) but, upon over-expression of *lncTAM34a*, increases to detectable levels, we hypothesized that *lncTAM34a* is capable of regulating *miR34a* expression via a transcriptional mechanism. To ascertain if this is actually the case, we performed chromatin immunoprecipitation (ChIP) for phosphorylated polymerase II (polII) at the *miR34a* HG promoter in both *lncTAM34a* overexpressing and mock control cell lines. Our results indicated a clear increase in phosphorylated polII binding at the *miR34a* promoter upon *lncTAM34a* over-expression indicating the ability of *lncTAM34a* to transcriptionally regulate *miR34a* levels (**Fig. 3d**).

### Low *lncTAM34a* expression levels are associated with decreased survival

As *TP53* mutations and low expression of *miR34a* have been associated with worse prognosis in cancer, we compared survival rates of samples with low expression of *lncTAM34a* (bottom 10th percentile) to control samples in 17 cancer types from TCGA (**Figure 4-Supplement 1**) (Gallardo et al. 2009, Zenz et al. 2009, Liu et al. 2011). To correct for the effect of *TP53* mutations we focused on non-TP53 mutated samples, and noted a worse survival for the low expression group in several cancers. This effect was most pronounced in papillary kidney cancer (unadjusted *P*=0.00095; **Fig. 4a**). By systematically comparing 5-year survival probabilities between the low expression group and the control group for each cancer we found a median reduction of 5-year survival probability of 9.6% (*P*=0.083; Wilcoxon signed rank test; **Fig. 4b**). Furthermore, we found that *lncTAM34a* expression showed similar patterns in terms of direction and strength of association with 5-year survival probability as *miR34a* expression (r=0.57, *P*=0.037) and TP53 mutations (r=0.80, *P*=0.00054) across the different cancer types (**Fig. 4b**). Although these results do not implicate any causal relationship, they do indicate a striking similarity between the association of worse prognosis and *TP53* mutations, low *miR34a*, and low *lncTAM34a* expression.

**Figure 4:**
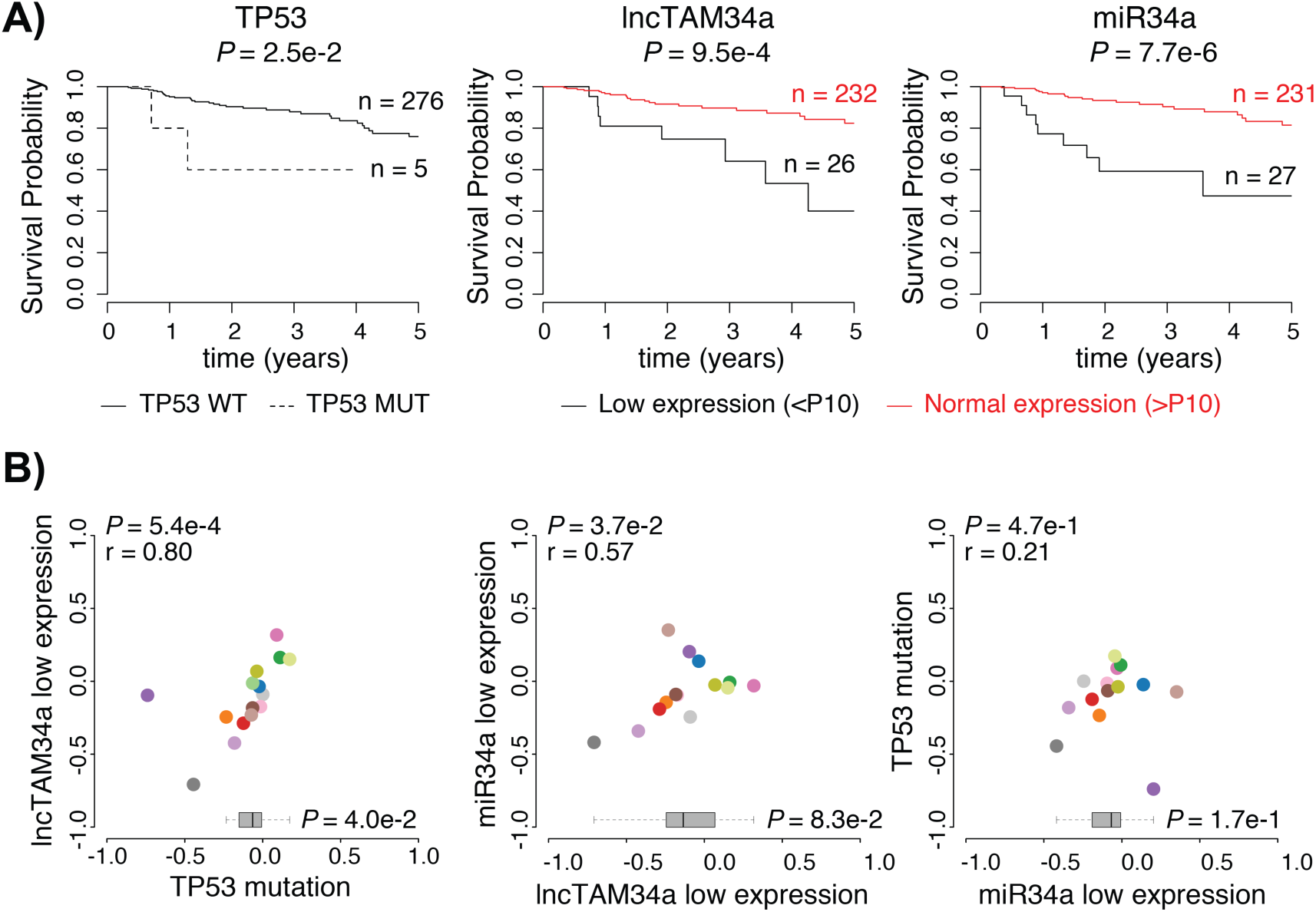
Survival analysis in TCGA cancers. **A)** Kaplan-Meier survival curves comparing the effects of TP53-mutated samples (left), low *lncTAM34a* expression (middle) and low *miR34a* expression (right) to control samples in papillary kidney cancer (results for other cancers in **Figure 4-Supplement 1**). Middle and Right panel include only TP53 wild type patients where RNAseq data exists. B) Correlation analysis between the effects on the 5-year survival probability of *TP* 53-mutated samples, low *lncTAM34a* expression, and low *miR34a* expression as indicated. For each variable the 5-year survival probability was compared to the control group (negative values indicate lower survival, positive values indicate higher survival). Spearman correlation coefficients are given on the top left of each plot. Each dot indicates one cancer type (see Fig. 1c for legend). Boxplots on the bottom summarize the effects for the parameter on the x-axis, with indication of *P* values, as calculated using paired Wilcoxon signed rank test. Low expression was defined as *TP53* non-mutated samples having expression values in the bottom 10th percentile.

**Figure 5:**
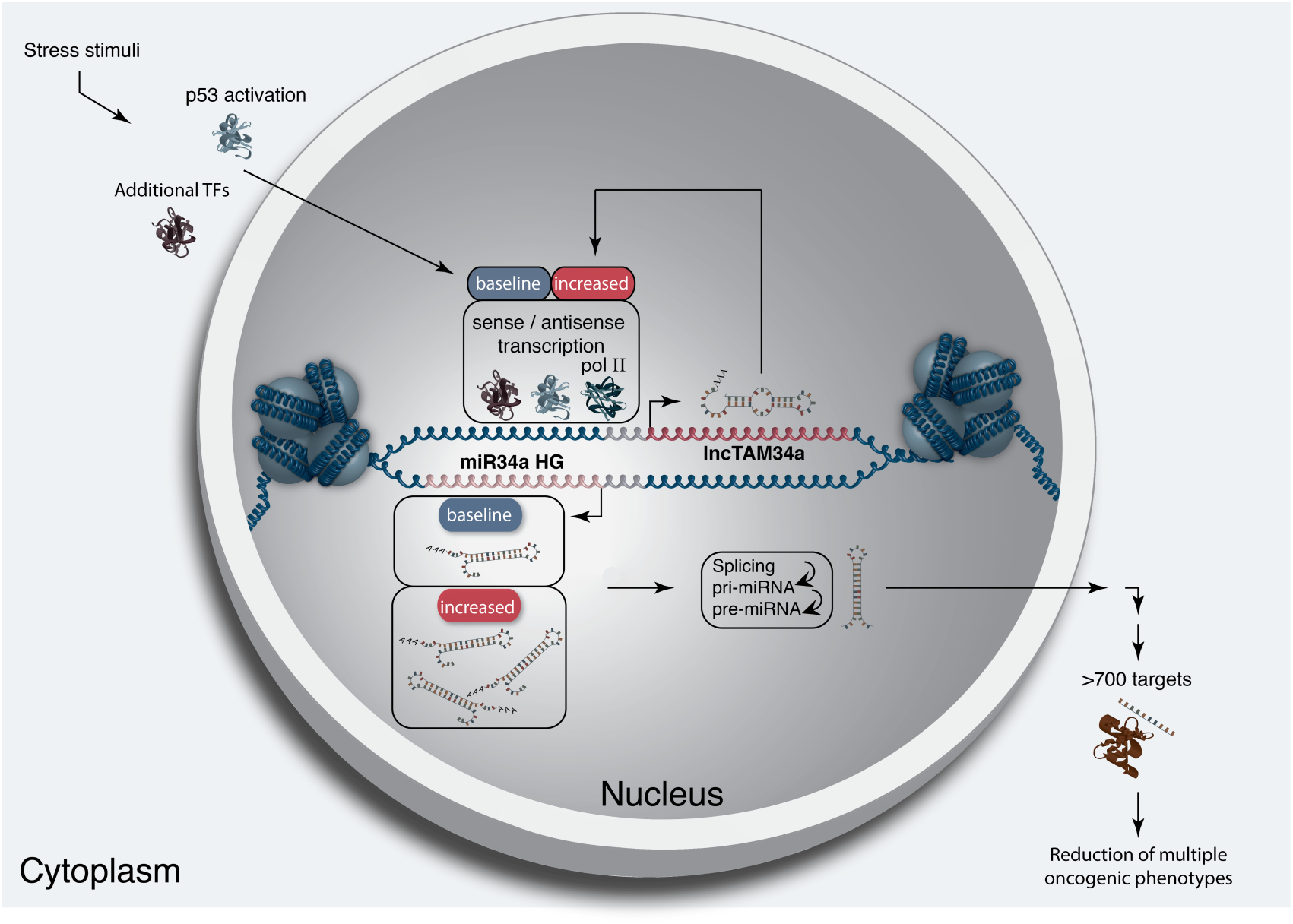
A graphical summary of the proposed *lncTAM34a* function. Stress stimuli, originating in the cytoplasm or nucleus, activate TP53 as well as additional factors. These factors then bind to the *miR34a* promoter and drive baseline transcription levels of the sense and antisense strands. *lncTAM34a* serves to further increase *miR34a* HG transcription levels resulting in enrichment of polymerase II at the *miR34a* promoter and a positive feed-forward loop. *miR34a* HG then, in turn, is spliced and processed in multiple steps before the mature *miR34a* binds to the RISC complex allowing it to repress its targets and exert its tumor suppressive effects.

## Discussion

Multiple studies have previously shown asRNAs to be crucial for the appropriate regulation of cancer-associated protein-coding genes and that their dysregulation can lead to perturbance of tumor suppressive and oncogenic pathways, as well as, cancer-related phenotypes (Yu et al. 2008, Yap et al. 2010, Serviss et al. 2014, Balbin et al. 2015). Here we show that asRNAs are also capable of regulating cancer-associated miRNAs resulting in similar consequences as protein-coding gene dysregulation (**Fig. 4**). Interestingly, we show that, both in the presence and absence of *TP53, lncTAM34a* provides an additional regulatory level to control *miR34a* expression in both homeostasis and upon encountering various forms of cellular stress. Furthermore, we find that *lncTAM34a-mediated* increase in *miR34a* expression is sufficient to drive the appropriate cellular responses to these stress stimuli (**Fig. 2d and Fig. 3c**). Previous studies have exploited various molecular biology methods to up-regulate *miR34a* expression in cells lacking wild type *TP53* (Liu et al. 2011, Ahn et al. 2012, Yang et al. 2012, Stahlhut et al. 2015, Wang et al. 2015). In this study, we demonstrate a novel, endogenous mechanism of *miR34a* regulation that has similar phenotypic outcomes as has been previously shown for *miR34a* induction in a *TP53* deficient background.

In agreement with previous studies, we demonstrate that upon encountering various types of cellular stress, TP53 in concert with additional factors initiates transcription at the *miR34a* locus, thus increasing the levels of *lncTAM34a* and *miR34a* (Rashi-Elkeles et al. 2014, Hunten et al. 2015, Leveille et al. 2015, Ashouri et al. 2016, Kim et al. 2017). We found that over-expression of *lncTAM34a* leads to recruitment of polII to the *miR34a* promoter and hypothesize that *lncTAM34a* may provide positive feedback for *miR34a* expression whereby it serves as a scaffold for the recruitment of additional factors that facilitate polII-mediated transcription. In this manner, *miR34a* expression is induced, driving a shift towards a reduction in growth factor signaling, senescence, and in some cases apoptosis. On the other hand, in cells without functional TP53, other factors, which typically act independently or in concert with TP53, may initiate transcription of the *miR34a* locus. Due to the fact that *lncTAM34a* can alter *miR34a* expression in these cells, we suggest that it is interacting with one of these additional factors, possibly recruiting it to the *miR34a* locus in order to drive *miR34a* transcription, similar to mechanisms described for other lncRNAs (Hung et al. 2011, Ng et al. 2012, Ng et al. 2013). The head-to-head orientation of the *miR34a* HG and *lncTAM34a* causes sequence complementarity between the RNA and the promoter DNA, making targeting by direct binding an attractive mechanism. Previous reports have also illustrated the ability of asRNAs to form hybrid DNA:RNA R-loops and, thus, facilitate an open chromatin structure and the transcription of the sense gene (Boque-Sastre et al. 2015). The fact that the p1 construct only contains a small portion (~300 bp) of the *lncTAM34a* transcript indicates that this portion is sufficient to give rise to at least a partial *miR34a* inducing response and therefore, that *lncTAM34a* may be able to facilitate *miR34a* expression independent of additional factors (**Fig 2d, Figure 2-Figure Supplement 2a**). Nevertheless, further work will need to be performed to explore the mechanism whereby *lncTAM34a* regulates *miR34a* gene expression.

An antisense transcript arising from the *miR34a* locus, *Lnc34a,* has been previously reported to negatively regulate the expression of *miR34a* (Wang et al. 2016). Although the *Lnc34a* and *lncTAM34a* transcripts share some sequence similarity, we believe them to be separate RNAs that are, potentially, different isoforms of the same gene. We utilized CAGE and RNAseq data from the ENCODE project to evaluate the presence of *lncTAM34a* and *Lnc34a* in 28 and 36 commonly used cancer cell lines, respectively. Although the results show the presence of *lncTAM34a* in these cell lines, we find no evidence for *Lnc34a* transcription (**Supplementary Document 1**). These results are in line with the findings of Wang et al. indicating that *Lnc34a* is highly expressed in colon cancer stem cell spheres compared to all other cell types used in their study and may not be broadly expressed in other tissues or tumor types. The fact that *lncTAM34a* and *Lnc34a* would appear to have opposing roles in their regulation of *miR34a*, further underlines the complexity of the regulation at this locus.

Clinical trials utilizing *miR34a* replacement therapy have previously been conducted but, disappointingly, were terminated after adverse side effects of an immunological nature were observed in several of the patients (Slabakova et al. 2017). Although it is not presently clear if these side effects were caused by *miR34a* or the liposomal carrier used to deliver the miRNA, the multitude of evidence indicating *miR34a*’s crucial role in oncogenesis still makes its therapeutic induction an interesting strategy and needs further investigation.

Our results indicate an association between survival probability and low *lncTAM34a* expression making it an attractive candidate for controlled preclinical studies. Due to the *lncTAM34a-mediated* positive feedback on *miR34a* expression, initiation of this feedback mechanism may provide a sustained *miR34a* induction in a relatively more robust manner than *miR34a* replacement alone. In summary, our results have identified *lncTAM34a* as a vital component in the regulation of *miR34a* and its particular importance in typical examples of cellular stress encountered in cancer. On a broader level, the conclusions drawn in this study provide an example of asRNA-mediated regulation of a clinically relevant cancer-associated miRNA and contribute to fundamental knowledge concerning *miR34a* regulation.

## Materials and Methods

### Cell Culture

All cell lines were cultured at 5% CO_2_ and 37°C with HEK293T, Saos2, and Skov3 cells cultured in DMEM high glucose (GE Healthcare Life Sciences, Hyclone, Amersham. UK, Cat# SH30081), HCT116 and U2OS cells in McCoy’s 5a (ThermoFisher Scientific, Pittsburgh, MA, USA. Cat# SH30200), and PC3 cells in RPMI (GE Healthcare Life Sciences, Hyclone, Cat# SH3009602) and 2 mM L-glutamine (GE Healthcare Life Sciences, Hyclone, Cat# SH3003402). All growth mediums were supplemented with 10% heat-inactivated FBS (ThermoFisher Scientific, Gibco, Cat# 12657029) and 50 pg/ml of streptomycin (ThermoFisher Scientific, Gibco, Cat# 15140122) and 50 μg/ml of penicillin (ThermoFisher Scientific, Gibco, Cat# 15140122). All cell lines were purchased from ATCC, tested negative for mycoplasma, and their identity was verified via STR profiling.

### Bioinformatics, Data Availability, and Statistical Testing

The USCS genome browser (Kent et al. 2002) was utilized for the bioinformatic evaluation of antisense transcription utilizing the RefSeq (O’Leary et al. 2016) gene annotation track.

All raw experimental data, code used for analysis, and supplementary methods are available for review at (Serviss 2017) and are provided as an R package. All analysis took place using the R statistical programming language (Team 2017) using external packages that are documented in the package associated with this article (Wilkins, Chang 2014, Wickham 2014, Therneau 2015, Wickham 2016, Allaire et al. 2017, Arnold 2017, Wickham 2017, Wickham 2017, Wickham 2017, Xiao 2017, Xie 2017). The package facilitates replication of the operating system and package versions used for the original analysis, reproduction of each individual figure and figure supplement included in the article, and easy review of the code used for all steps of the analysis, from raw-data to figure.

The significance threshold (alpha) in this study was set to 0.05. Statistical testing was performed using an unpaired two sample Student’s t-test unless otherwise specified.

### Coding Potential

Protein-coding capacity was evaluated using the Coding-potential Assessment Tool (Wang et al. 2013) and Coding-potential Calculator (Kong et al. 2007) with default settings. Transcript sequences for use with Coding-potential Assessment Tool were downloaded from the UCSC genome browser using the Ensembl accessions: *HOTAIR* (ENST00000455246), *XIST* (ENST00000429829), β-actin (ENST00000331789), Tubulin (ENST00000427480), and *MYC* (ENST00000377970). Transcript sequences for use with Coding-potential Calculator were downloaded from the UCSC genome browser using the following IDs: *HOTAIR* (uc031qho.1), β-actin (uc003soq.4).

### Peptide identification in MS/MS spectra

Orbitrap raw MS/MS files for 11 human cell lines were downloaded from the PRIDE repository (PXD002395; (Geiger et al. 2012)) converted to mzML format using msConvert from the ProteoWizard tool suite (Holman et al. 2014). Spectra were then searched using MSGF+ (v10072) (Kim et al. 2014) and Percolator (v2.08) (Granholm et al. 2014). All searches were done against the human protein subset of Ensembl 75 in the Galaxy platform (Boekel et al. 2015) supplemented with the 6 frame translation of both the annotated (LOC102724571; hg38) and PCR cloned sequence of *lncTAM34a* (supplementary data; (Serviss 2017)). MSGF+ settings included precursor mass tolerance of 10 ppm, fully-tryptic peptides, maximum peptide length of 50 amino acids and a maximum charge of 6. Fixed modification was carbamidomethylation on cysteine residues; a variable modification was used for oxidation on methionine residues. Peptide Spectral Matches found at 1% FDR (false discovery rate) were used to infer peptide identities. The output from all searches are available in (Serviss 2017).

### shRNAs

shRNA-expressing constructs were cloned into the U6M2 construct using the BglII and KpnI restriction sites as previously described (Amarzguioui et al. 2005). shRNA constructs were transfected using Lipofectamine 2000 or 3000 (ThermoFisher Scientific, Cat# 12566014 and L3000015). The sequences targeting renilla is as follows: shRenilla 1.1 (AAT ACA CCG CGC TAC TGG C), shRenilla 2.1 (TAA CGG GAT TTC ACG AGG C).

### Bi-directional Promoter Cloning

The overlapping region (p1) corresponds with the sequence previously published as the TP53 binding site in (Raver-Shapira et al. 2007) which we synthesized, cloned into the pLucRluc construct (Polson et al. 2011), and sequenced to verify its identity.

### Promoter Activity

Cells were co-transfected with the p1 renilla/firefly bidirectional promoter construct (Polson et al. 2011) and GFP by using Lipofectamine 2000 (Life Technologies, Cat# 12566014). The expression of GFP and luminescence was measured 24 h post transfection by using the Dual-Glo Luciferase Assay System (Promega, Cat# E2920) and detected by the GloMax-Multi+ Detection System (Promega, Cat# SA3030). The expression of luminescence was normalized to GFP.

### Generation of U6-expressed *lncTAM34a* Lentiviral Constructs

The U6 promoter was amplified from the U6M2 cloning plasmid (Amarzguioui et al. 2005) and ligated into the Not1 restriction site of the pHIV7-IMPDH2 vector (Turner et al. 2012). *lncTAM34a* was PCR amplified and subsequently cloned into the Nhe1 and Pac1 restriction sites in the pHIV7-IMPDH2-U6 plasmid.

### Lentiviral Particle production, infection, and selection

Lentivirus production was performed as previously described in (Turner et al. 2012). Briefly, HEK293T cells were transfected with viral and expression constructs using Lipofectamine 2000 (ThermoFisher Scientific, Cat# 12566014), after which viral supernatants were harvested 48 and 72 hours post-transfection. Viral particles were concentrated using PEG-IT solution (Systems Biosciences, Palo Alto, CA, USA. Cat# LV825A-1) according to the manufacturer’s recommendations. HEK293T cells were used for virus titration and GFP expression was evaluated 72hrs post-infection via flow cytometry (LSRII, BD Biosciences, San Jose, CA, USA) after which TU/ml was calculated.

Stable lines were generated by infecting cells with a multiplicity of infection of 1 and subsequently initiating 1-2 μM mycophenolic acid-based (Merck, Kenilworth, NJ, USA. Cat# M5255) selection 48-72 hours post-infection. Cells were expanded as the selection process was monitored via flow cytometry analysis (LSRII, BD Biosciences) of GFP and selection was terminated once > 90% of the cells were GFP positive. Quantification of *lncTAM34a* overexpression and *miR34a* was performed in biological quintuplet for all cell lines.

### Western Blotting

Samples were lysed in 50 mM Tris-HCl (Sigma Aldrich, St. Louis, MO, USA. Cat# T2663), pH 7.4, 1% NP-40 (Sigma Aldrich, Cat# I8896), 150 mM NaCl (Sigma Aldrich, Cat# S5886), 1 mM EDTA (Promega, Madison, WI, USA. Cat# V4231), 1% glycerol (Sigma Aldrich, Cat# G5516), 100 μM vanadate (Sigma Aldrich, Cat# S6508), protease inhibitor cocktail (Roche Diagnostics, Basel, Switzerland, Cat# 004693159001) and PhosSTOP (Roche Diagnostics, Cat# 04906837001). Lysates were subjected to SDS-PAGE and transferred to PVDF membranes. The proteins were detected by western blot analysis by using an enhanced chemiluminescence system (Western Lightning-ECL, PerkinElmer, Waltham, MA, USA. Cat# NEL103001EA). Antibodies used were specific for CCND1 1:1000 (Cell Signaling, Danvers, MA, USA. Cat# 2926), and GAPDH 1:5000 (Abcam, Cambridge, UK, Cat# ab9485). All western blot quantifications were performed using ImageJ (Schneider et al. 2012).

### RNA Extraction and cDNA Synthesis

For downstream SYBR green applications, RNA was extracted using the RNeasy mini kit (Qiagen, Venlo, Netherlands, Cat# 74106) and subsequently treated with DNase (Ambion Turbo DNA-free, ThermoFisher Scientific, Cat# AM1907). 500ng RNA was used for cDNA synthesis using MuMLV (ThermoFisher Scientific, Cat# 28025013) and a 1:1 mix of oligo(dT) and random nanomers.

For analysis of miRNA expression with Taqman, samples were isolated with TRIzol reagent (ThermoFisher Scientific, Cat# 15596018) and further processed with the miRNeasy kit (Qiagen, Cat# 74106). cDNA synthesis was performed using the TaqMan MicroRNA Reverse Transcription Kit (ThermoFisher Scientific, Cat# 4366597) using the corresponding oligos according to the manufacturer’s recommendations.

### QPCR and PCR

PCR was performed using the KAPA2G Fast HotStart ReadyMix PCR Kit (Kapa Biosystems, Wilmington, MA, USA, Cat# KK5601) with corresponding primers. QPCR was carried out using KAPA 2G SYBRGreen (Kapa Biosystems, Cat# KK4602) using the Applied Biosystems 7900HT machine with the cycling conditions: 95 °C for 3 min, 95 °C for 3 s, 60 °C for 30 s.

QPCR for miRNA expression analysis was performed according to the primer probe set manufacturers recommendations (ThermoFisher Scientific) and using the TaqMan Universal PCR Master Mix (ThermoFisher Scientific, Cat# 4304437) with the same cycling scheme as above. Primer and probe sets for TaqMan were also purchased from ThermoFisher Scientific (Life Technologies at time of purchase, TaqMan^®^ MicroRNA Assay, hsa-miR-34a, human, Cat# 4440887, Assay ID: 000426 and Control miRNA Assay, RNU48, human, Cat# 4440887, Assay ID: 001006).

The ΔΔCt method was used to quantify gene expression. All QPCR-based experiments were performed in at least technical duplicate. Primers for all PCR-based experiments are listed in **Supplementary Document 2** and arranged by figure.

### Cell Cycle Distribution

Cells were washed in PBS and fixed in 4% paraformaldehyde at room temperature overnight. Paraformaldehyde was removed, and cells were re-suspended in 95% EtOH. The samples were then rehydrated in distilled water, stained with DAPI and analyzed by flow cytometry on a LSRII (BD Biosciences) machine. Resulting cell cycle phases were quantified using the ModFit software (Verity Software House, Topsham, ME, USA). Experiments were performed in biological quadruplet (PC3) or triplicate (Skov3). The log2 fraction of cell cycle phase was calculated for each replicate and a two sample t-test was utilized for statistical testing.

### 3’ Rapid Amplification of cDNA Ends

3’-RACE was performed as described as previously in (Johnsson et al. 2013). Briefly, U2OS cell RNA was polyA-tailed using yeast polyA polymerase (ThermoFisher Scientific, Cat# 74225Z25KU) after which cDNA was synthesized using oligo(dT) primers. Nested-PCR was performed first using a forward primer in *lncTAM34a* exon 1 and a tailed oligo(dT) primer followed by a second PCR using an alternate *lncTAM34a* exon 1 primer and a reverse primer binding to the tail of the previously used oligo(dT) primer. PCR products were gel purified and cloned the Strata Clone Kit (Agilent Technologies, Santa Clara, CA, USA. Cat# 240205), and sequenced.

### Chromatin Immunoprecipitation

The ChIP was performed as previously described in (Johnsson et al. 2013) with the following modifications. Cells were crosslinked in 1% formaldehyde (Merck, Cat# 1040039025), quenched with 0.125M glycine (Sigma Aldrich, Cat# G7126), and lysed in cell lysis buffer comprised of: 5mM PIPES (Sigma Aldrich, Cat# 80635), 85mM KCL (Merck, Cat# 4936), 0.5% NP40 (Sigma Aldrich, Cat# I8896), protease inhibitor (Roche Diagnostics, Cat# 004693159001). Samples were then sonicated in 50mM TRIS-HCL pH 8.0 (Sigma Aldrich, MO, USA, Cat# T2663) 10mM EDTA (Promega, WI, USA, Cat# V4231), 1% SDS (ThermoFisher Scientific, Cat# AM9822), and protease inhibitor (Roche Diagnostics, Cat# 004693159001) using a Bioruptor Sonicator (Diagenode, Denville, NJ, USA). Samples were incubated over night at 4°C with the polII antibody (Abcam, Cat# ab5095) and subsequently pulled down with Salmon Sperm DNA/Protein A Agarose (Millipore, Cat# 16157) beads. DNA was eluted in an elution buffer of 1% SDS (ThermoFisher Scientific, Cat# AM9822) 100mM NaHCO3 (Sigma Aldrich, Cat# 71631), followed by reverse crosslinking, RNaseA (ThermoFisher Scientific, Cat# 1692412) and protease K (New England Biolabs, Ipswich, MA, USA, Cat# P8107S) treatment. The DNA was eluted using Qiagen PCR purification kit (Cat# 28106) and quantified via QPCR. QPCR was performed in technical duplicate using the standard curve method and reported absolute values. The fraction of input was subsequently calculated using the mean of the technical replicates followed by calculating the fold over the control condition. Statistical testing was performed using 4 biological replicates with the null hypothesis that the true log2 fold change values were equal to zero.

### Confluency Analysis

Cells were incubated in the Spark Multimode Microplate (Tecan, Männedorf, Switzerland) reader for 48 hours at 37°C with 5% CO_2_ in a humidity chamber in either normal medium or HBSS (ThermoFisher Scientific, Cat# 14025092). Confluency was measured every hour using bright-field microscopy and the percentage of confluency was reported via the plate reader’s inbuilt algorithm. Percentage of confluency was normalized to the control sample in each condition (shown in figure) and then ranked to move the data to a linear scale. Using the mean of the technical duplicates in three biological replicates, the rank was then use to construct a linear model, of the dependency of the rank on the time and cell lines variables for each growth condition. Reported *P* values are derived from the t-test, testing the null hypothesis that the coefficient estimate of the cell line variable is equal to 0.

### Pharmacological Compounds

Doxorubicin was purchased from Teva (Petah Tikva, Israel, cat. nr. 021361).

### Cellular Localization Analysis

Quantified RNAseq data from 11 cell lines from the GRCh38 assembly was downloaded from the ENCODE project database and quantifications for *lncTAM34a* (ENSG00000234546), GAPDH (ENSG00000111640), and MALAT1 (ENSG00000251562) were extracted. Cell lines for which data was downloaded include: A549, GM12878, HeLa-S3, HepG2, HT1080, K562 MCF-7, NCI-H460, SK-MEL-5, SK-N-DZ, SK-N-SH. Initial exploratory analysis revealed that several cell lines should be removed from the analysis due to a) a larger proportion of GAPDH in the nucleus than cytoplasm or b) variation of *lncTAM34a* expression is too large to draw conclusions, or c) they have no or low (<6 TPM) *lncTAM34a* expression. Furthermore, only polyadenylated libraries were used in the final analysis, due to the fact that the cellular compartment enrichment was improved in these samples. All analyzed genes are reported to be polyadenylated. In addition, only samples with 2 biological replicates were retained. For each cell type, gene, and biological replicate the fraction of transcripts per million (TPM) in each cellular compartment was calculated as the fraction of TPM in the specific compartment by the total TPM. The mean and standard deviation for the fraction was subsequently calculated for each cell type and cellular compartment and this information was represented in the final figure.

### CAGE Analysis

All available CAGE data from the ENCODE project (Consortium 2012) for 36 cell lines was downloaded from the UCSC genome browser (Kent et al. 2002) for genome version hg19. Of these, 28 cell lines had CAGE transcription start sites (TSS) mapping to the plus strand of chromosome 1 and in regions corresponding to 200 base pairs upstream of the *Lnc34a* start site (9241796 - 200) and 200 base pairs upstream of the GENCODE annotated *lncTAM34a* start site (9242263 + 200). These cell lines included: HFDPC, H1-hESC, HMEpC, HAoEC, HPIEpC, HSaVEC, GM12878, hMSC-BM, HUVEC, AG04450, hMSC-UC, IMR90, NHDF, SK-N-SH_RA, BJ, HOB, HPC-PL, HAoAF, NHEK, HVMF, HWP, MCF-7, HepG2, hMSC-AT, NHEM.f_M2, SkMC, NHEM_M2, and HCH. In total 74 samples were included. 17 samples were polyA-, 47 samples were polyA+, and 10 samples were total RNA. In addition, 34 samples were whole cell, 15 enriched for the cytosolic fraction, 15 enriched for the nucleolus, and 15 enriched for the nucleus. All CAGE transcription start sites were plotted and the RPKM of the individual reads was used to color each read to indicate their relative abundance. In cases where CAGE TSS spanned identical regions, the RPMKs of the regions were summed and represented as one CAGE TSS in the figure. In addition, a density plot shows the distribution of the CAGE reads in the specified interval.

### Splice Junction Analysis

All available whole cell (i.e. non-fractionated) spliced read data originating from the Cold Spring Harbor Lab in the ENCODE project (Consortium 2012) for 38 cell lines was downloaded from the UCSC genome browser (Kent et al. 2002). Of these cell lines, 36 had spliced reads mapping to the plus strand of chromosome 1 and in the region between the *Lnc34a* start (9241796) and transcription termination (9257102) site (note that *lncTAM34a* resides totally within this region). Splice junctions from the following cell lines were included in the final figure: A549, Ag04450, Bj, CD20, CD34 mobilized, Gm12878, H1hesc, Haoaf, Haoec, Hch, Helas3, Hepg2, Hfdpc, Hmec, Hmepc, Hmscat, Hmscbm, Hmscuc, Hob, Hpcpl, Hpiepc, Hsavec, Hsmm, Huvec, Hvmf, Hwp, Imr90, Mcf7, Monocd14, Nhdf, Nhek, Nhemfm2, Nhemm2, Nhlf, Skmc, and Sknsh. All splice junctions were included in the figure and colored according to the number of reads corresponding to each. In cases where identical reads were detected multiple times, the read count was summed and represented as one read in the figure.

### TCGA Data Analysis

RNAseq data and copy number data were downloaded from TCGA and processed as described previously (Ashouri et al. 2016). Briefly, RNAseq data were aligned to the human hg19 assembly and quantified using GENCODE (v19) annotated HTSeq-counts and FPKM normalizations. Expression data from *miR34a* and *lncTAM34a* (identified as RP3-510D11.2) were used for further analysis. Copy number amplitudes for GENCODE genes were determined from segmented copy-number data. Samples that were diploid for *lncTAM34a* were identified as those samples that had copy number amplitudes between −0.1 and 0.1.

Somatic mutation data were downloaded from the Genomics Data Commons data portal (GDC) as mutation annotation format (maf) files, called using Mutect2 on 30/10/2017 (v7) (Grossman et al. 2016).

Survival analysis was performed on TCGA vital state and follow-up data, downloaded from GDC on 27/10/2017 using the R survival package (Therneau 2015).

## Acknowledgments

The authors would like to kindly thank Martin Enge for his critical review of the manuscript and fruitful discussions. Dan Grandér, who played a significant role in the conceptualization and supervision of this project, sadly passed away before the initial submission of the manuscript. May he rest in peace.

**Figure Supplements**

Figure 1-Supplement 1: TCAG expression levels and correlation analysis statistics.

Figure 1-Supplement 2: Molecular characteristics of *lncTAM34a*.

Figure 2-Supplement 1: A schematic representation of the p1 construct.

Figure 2-Supplement 2: Evaluating the effects of *lncTAM34a* down-regulation.

Figure 3-Supplement 1: Physiological relevance of *lncTAM34a* over-expression.

Figure 3-Supplement 2: Effects of *lncTAM34a* over-expression on cyclin D1.

Figure 4-Supplement 1: Survival analysis in 17 cancers from TCGA.

Supplementary Document 1: Evaluating the relationship between *lncTAM34a* and *Lnc34a.*

Supplementary Document 2: A table of primers used in this study.

## Supplementary Figures

**Figure 1 Supplement 1:**
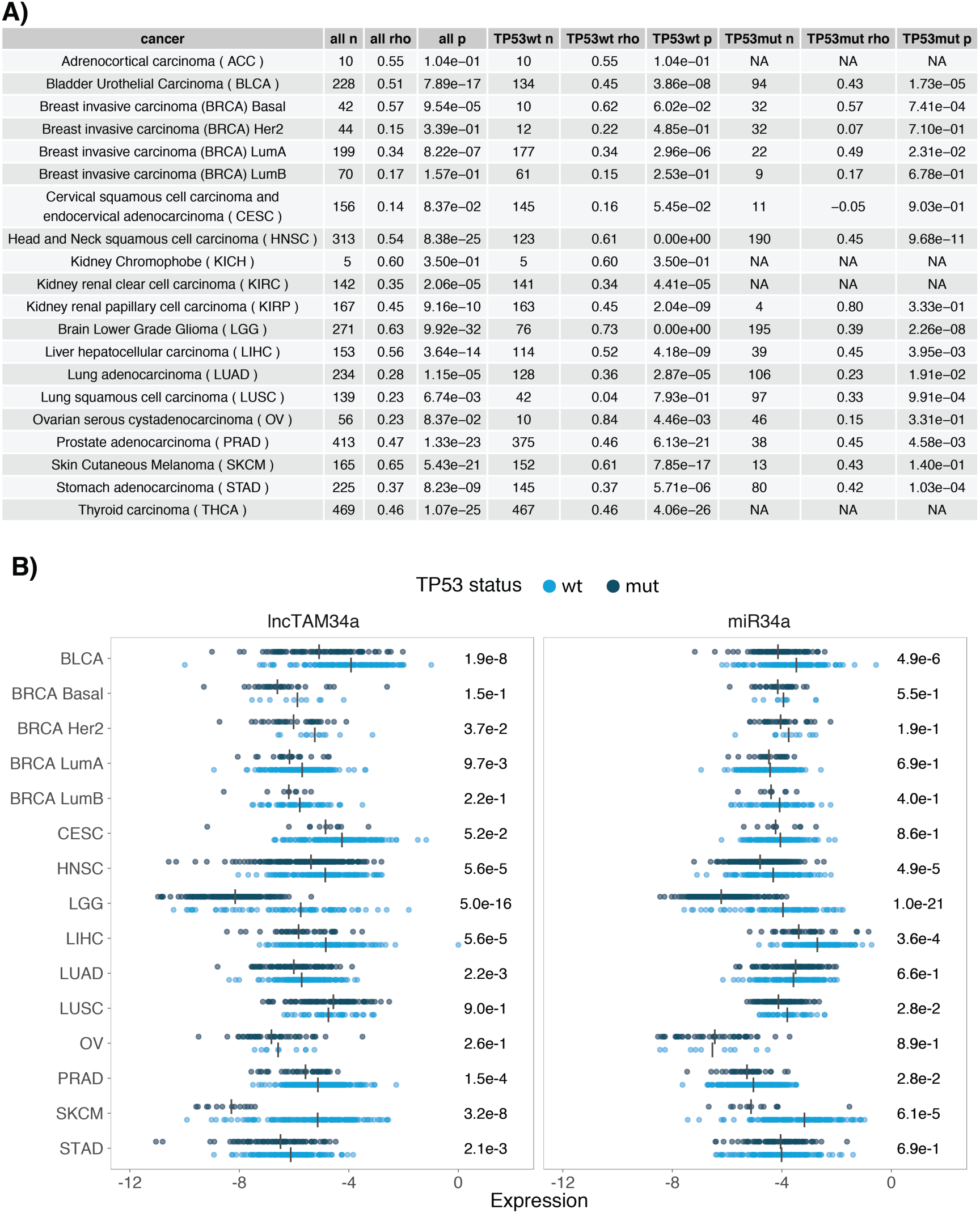
TCGA normalized expression levels and correlation analysis statistics. **A)** Spearman’s rho and *P* values (p) from the correlation analysis in Figure 1a between *miR34a* and *lncTAM34a* expression in *TP53* wild type (wt) and mutated (mut) samples within TCGA cancer types. NA indicates not applicable, due to a lack of data for the specific group. **B)** Expression levels of *miR34a* and *lncTAM34a* in *TP53* wt and nonsynonymous mutation samples. Expression was quantified by the log2 ratio of expression of the gene to its maximal expression value. Vertical lines indicate the median. *P* values are indicated on the right side of each panel and are derived from comparing the *TP53* wild type samples to the samples with a nonsynonymous mutation using a two-sided Wilcoxon signed rank test. Only cancers that had at least 5 samples per group were included. In addition, only samples that were diploid at the *miR34a* locus were used for the analysis to avoid copy number bias.

**Figure 1 Supplement 2:**
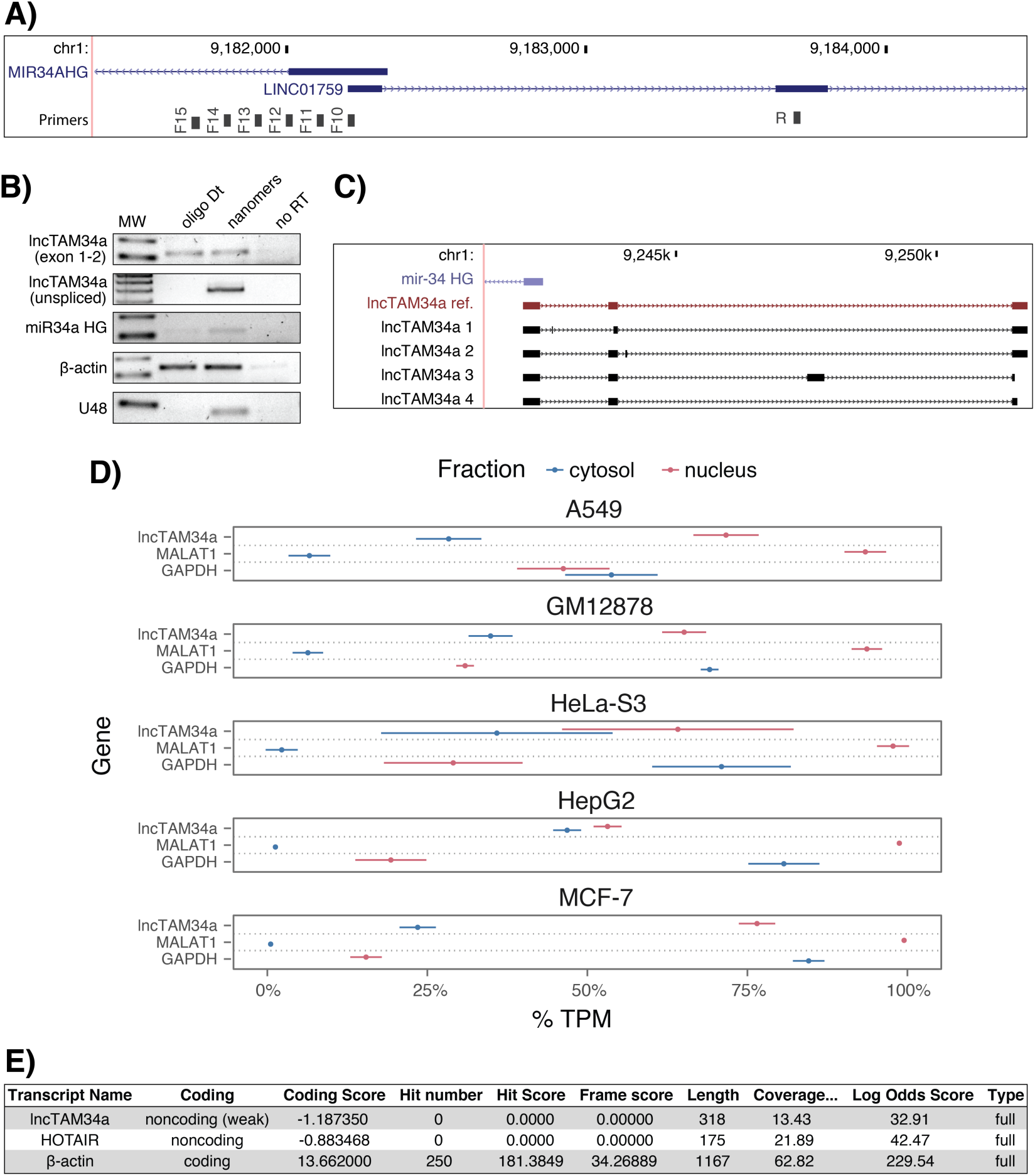
Molecular characteristics of *lncTAM34a*. **A)** A schematic representation of the primer placement in the primer walk assay. **B)** Polyadenylation status of spliced and unspliced *lncTAM34a* in HEK293T cells. **C)** Sequencing results from the analysis of *lncTAM34a* isoforms in U2OS cells. *lncTAM34a* ref. refers to the full-length transcript as defined by the 3’-RACE and the primer walk assay. **D)** Analysis of coding potential of the *lncTAM34a* transcript using the Coding-potential Calculator. **E)** RNAseq data from five fractionated cell lines in the ENCODE project showing the percentage of transcripts per million (TPM) for *lncTAM34a. MALAT1* (nuclear localization) and *GAPDH* (cytoplasmic localization) are included as fractionation controls. Points represent the mean and horizontal lines represent the standard deviation from two biological replicates.

**Figure 2 Supplement 1:**
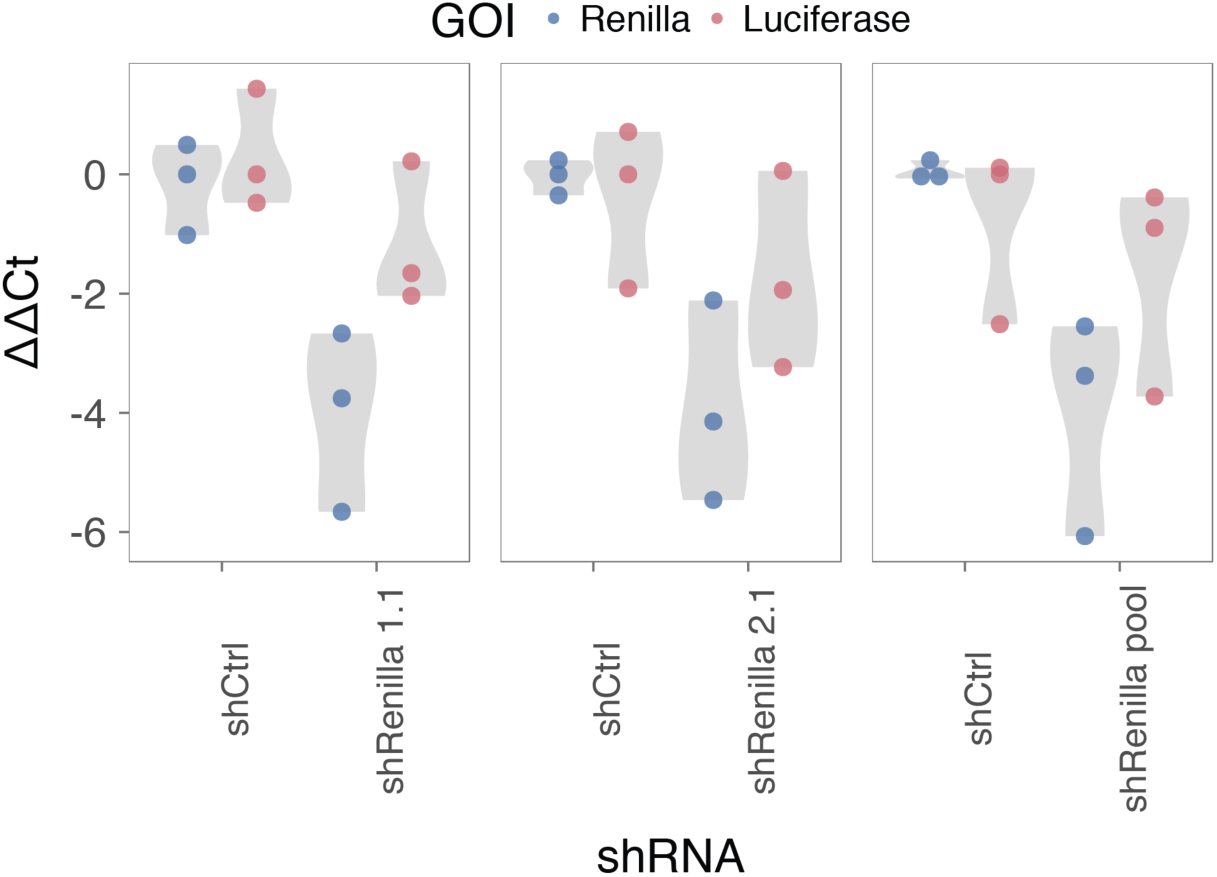
A schematic representation of the p1 construct. **A)** A UCSC genome browser illustration indicating the location of the promoter region cloned into the p1 construct including the conserved TP53-binding site. **B)** A representative picture of the p1 construct including forward (F) and reverse (R) primer locations and the renilla shRNA targeting site.

**Figure 2 Supplement 2:**
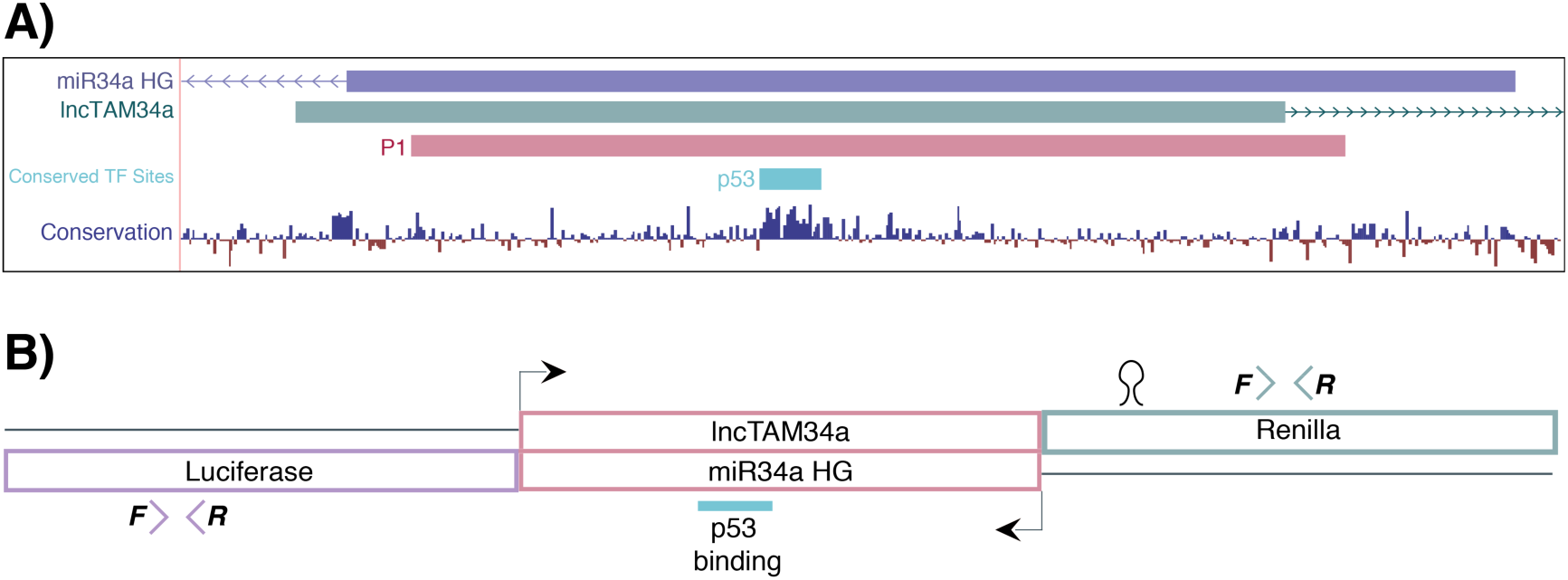
Evaluating the effects of *lncTAM34a* down-regulation. HEK293T cells were co-transfected with the p1 construct and either shRenilla or shControl. Renilla and luciferase levels were measured with QPCR 48 hours after transfection. Individual points represent independent experiments with the gray shadow indicating the density of the points. The experiment was performed in biological triplicate.

**Figure 3 Supplement 1:**
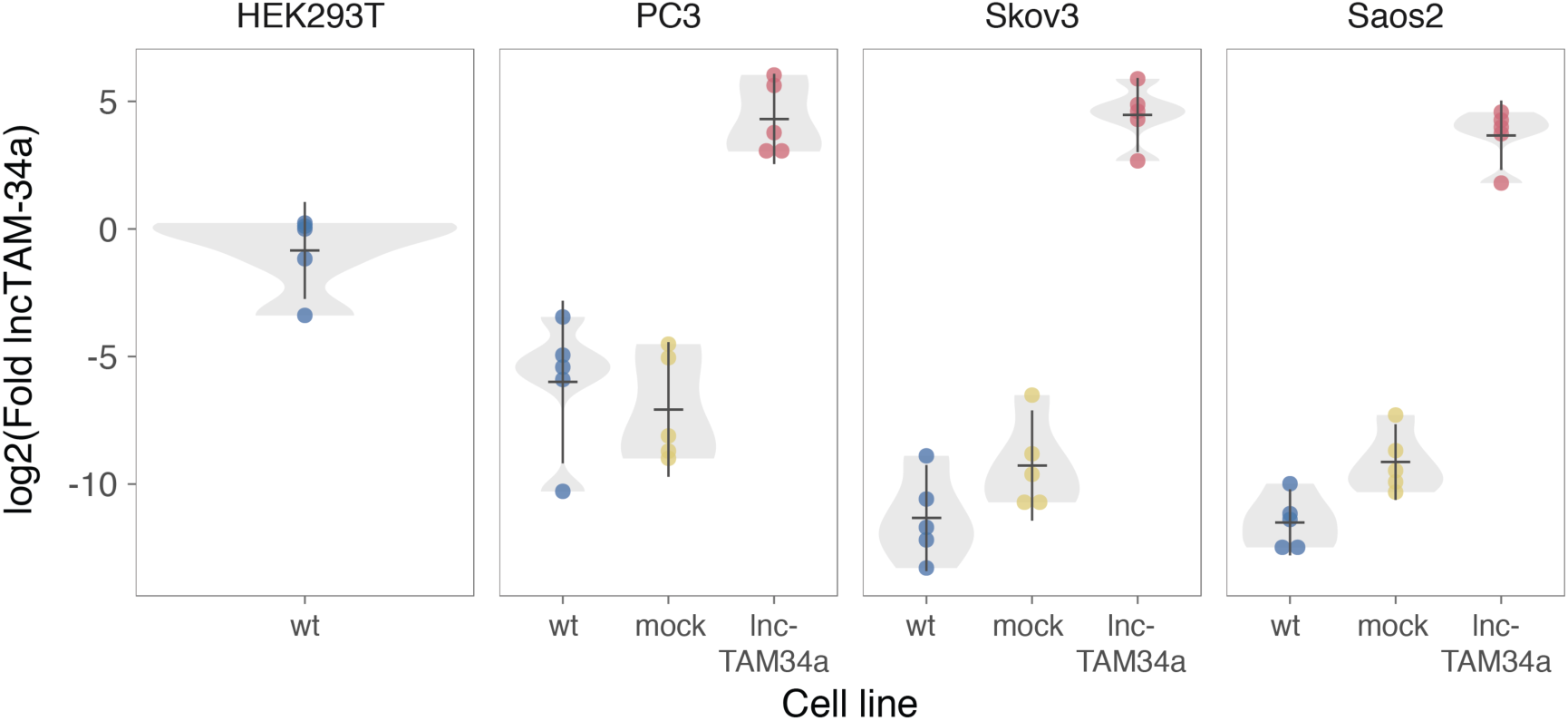
Physiological relevance of *lncTAM34a* over-expression. Comparison of *lncTAM34a* expression in HEK293T cells (high endogenous *lncTAM34a*), and the wild-type (wt), mock, and *lncTAM34a* over-expressing stable cell lines.

**Figure 3 Supplement 2:**
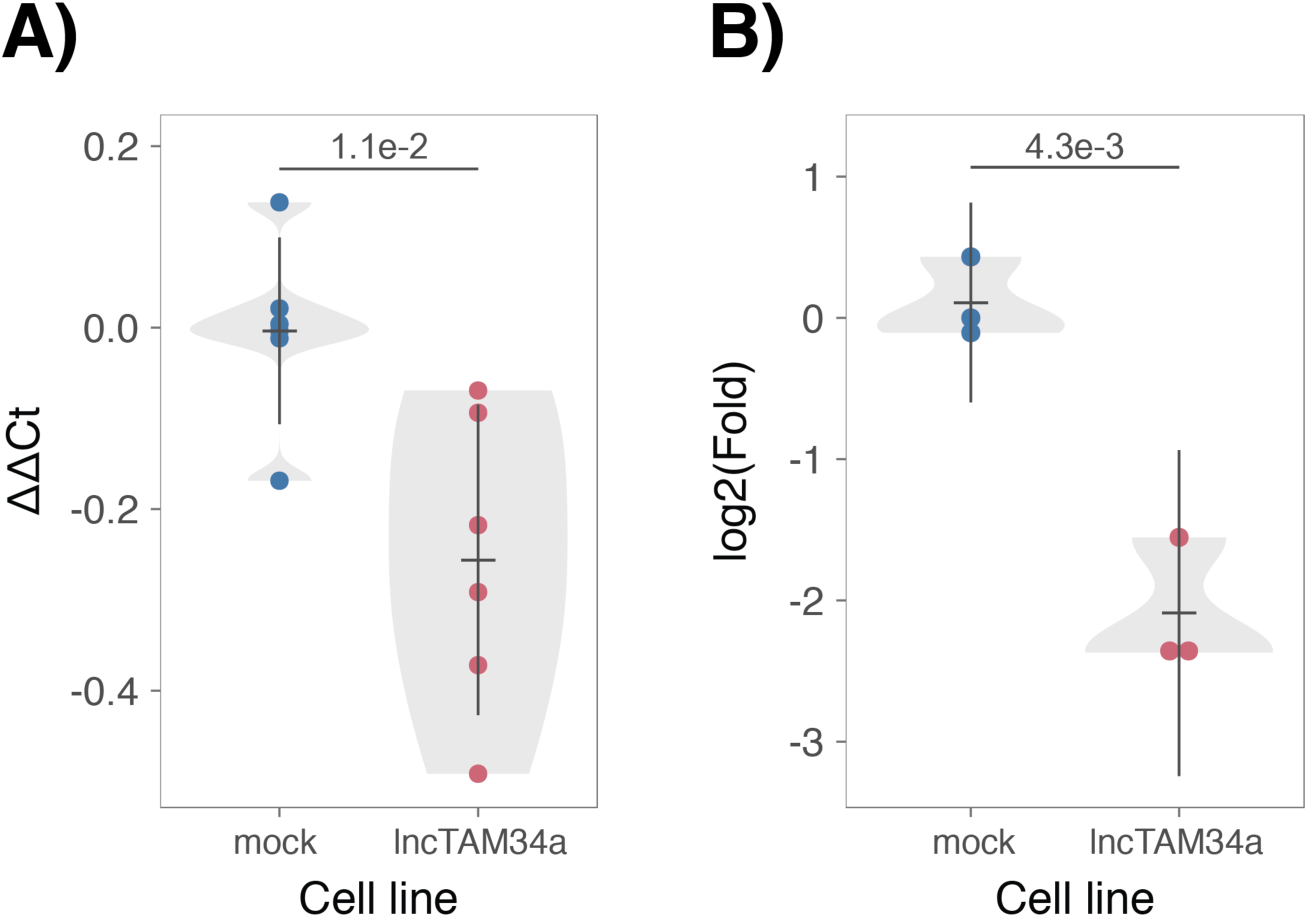
Effects of *lncTAM34a* over-expression on cyclin D1. CCND1 expression **(A)** and western blot quantification of protein levels **(B)** in *lncTAM34a* over-expressing PC3 stable cell lines. Experiments were performed in biological sextuplets **(A)** or triplicates **(B)**.

**Figure 4-Supplement 1:**
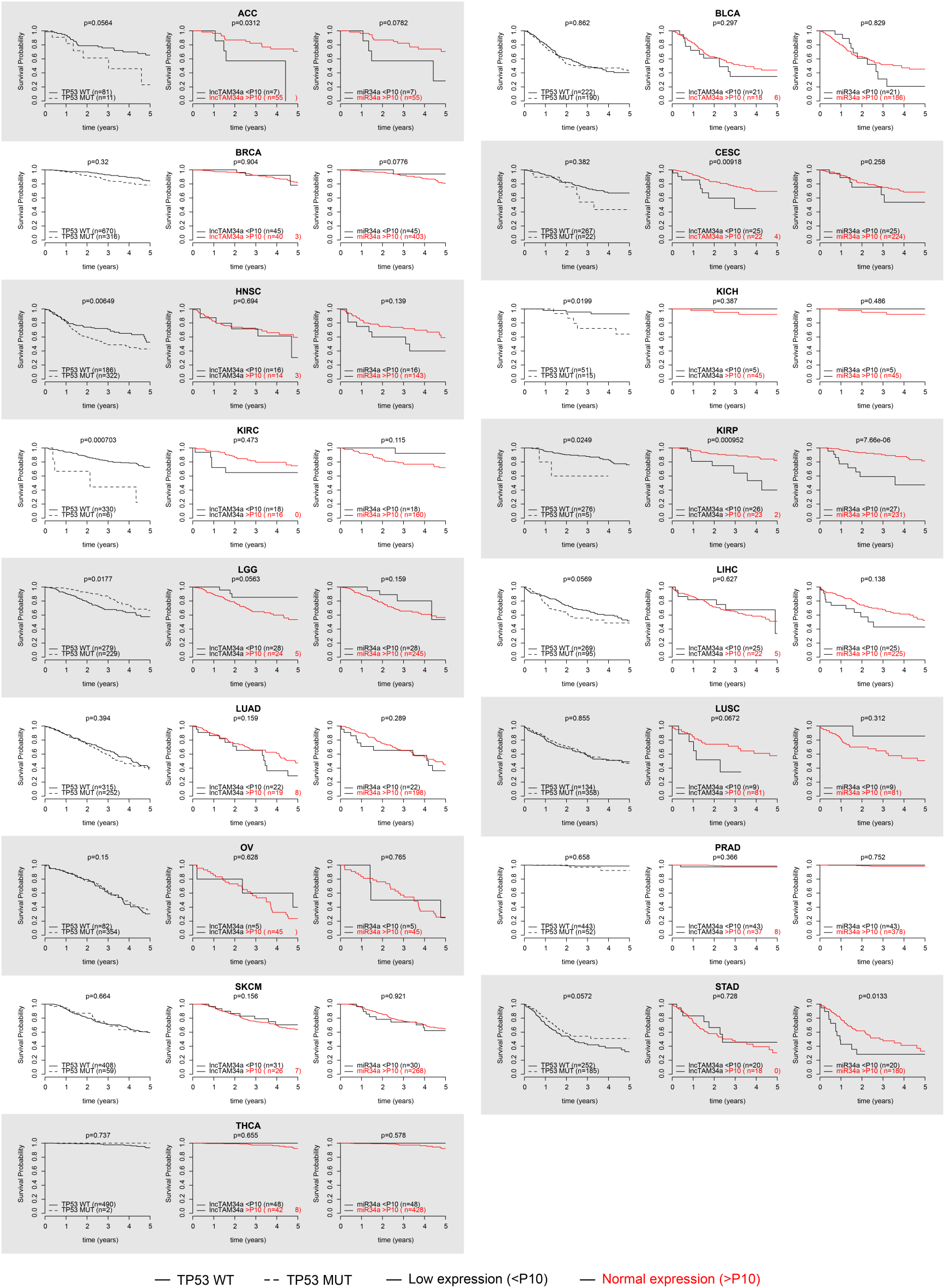
Survival analysis in 17 cancers from TCGA. Kaplan-Meier survival curves comparing the survival of TP53-mutated samples (left), low *lncTAM34a* expression (middle) and low *miR34a* expression (right) to control samples in 17 cancer types from TCGA. Low expression was defined as *TP53* non-mutated samples having expression values in the bottom 10th percentile.

